# Circadian modulation of spontaneous dopamine release shapes reward-evoked signaling in the nucleus accumbens

**DOI:** 10.64898/2026.04.22.720003

**Authors:** Jordan N. Cook, Mikael Gevorgyan, Jason Armitage, Jeff R. Jones

## Abstract

The circadian system is an important regulator of reward-related neural function and behavior. Dopamine (DA) release in the nucleus accumbens is a key component of reward processing, yet how circadian timing shapes DA release in relation to reward behavior remains unclear. Here, we investigated circadian rhythms in DA release and reward behavior using long-term fiber photometry paired with an automated reward delivery and measurement system. We found two distinct circadian rhythms in DA release: spontaneous DA, reflecting ongoing DA release not associated with reward, and reward-evoked DA, reflecting transient DA response during reward. Spontaneous DA peaked during the early subjective day, whereas reward-evoked peak DA peaked near the day-to-night transition. Both rhythms were distinct from reward behavior, which peaked during the early subjective night. Linear modeling further showed that the relationship between reward-evoked DA and reward behavior depended on circadian time, with greater DA responses occurring between late subjective day and early subjective night. Spontaneous baseline and reward-evoked DA were also negatively correlated, and this relationship was likewise modulated across circadian time. Together, these findings support a model in which circadian modulation of baseline DA may alter the gain of reward-evoked signaling, amplifying DA responses across behaviorally relevant times of day.

## INTRODUCTION

The brain’s reward system relies on circadian timing cues to generate daily rhythms in reward behavior [1,2]. In mammals, the motivation to engage in reward behavior is driven by the release of dopamine (DA) into the nucleus accumbens (NAc) [3,4]. Current literature suggests that striatal DA activity is regulated on a circadian timescale, including the synthesis, release, and reuptake of DA [5]. However, how circadian rhythms in DA release relate to rhythms in reward behavior remains unclear. This question has important implications for human health, as circadian dysfunction is comorbid with disorders affecting reward-related processes, including bipolar disorder, depression, and addiction [6,7]. Identifying circadian mechanisms that modulate DA release and reward behavior is therefore an important step towards understanding its broader role in neuropsychiatric disease pathology. Here, we investigate how circadian time modulates DA release in the NAc and its relationship to reward behavior.

The mammalian circadian system is organized hierarchically, with the suprachiasmatic nucleus (SCN) functioning as the central pacemaker [8,9]. A longstanding question in the field is how rhythmic information is encoded in brain regions outside the SCN. Nearly all nucleated cells express circadian genes, collectively known as the molecular clock. Across neuronal cell populations, these molecular clocks support local functions, including neural firing and neurotransmitter release [10–13]. Thus, one possible explanation for how rhythms in reward behavior are generated is that circadian information is encoded in local neural activity, such as the release of DA in the NAc.

DA release into the ventral striatum encodes many important properties of reward, such as novelty, value, and expectation. Novel rewards elicit greater DA neuron firing and release than expected ones, and this relationship also holds true for reward value [4,14,15]. Importantly, the dynamics of midbrain dopaminergic neuron firing during reward are partially dissociable from properties of DA release [16]. During fully predicted rewards, a measurable increase in accumbal DA still occurs despite there being no increase in the firing frequency of DAergic neurons. Furthermore, DA release in the NAc is directly involved in specific properties of motivation, such as action initiation and approach [17]. Thus, DA release, as opposed to just DAergic neuron activity, is a particularly important neural correlate for reward behavior.

Despite this, technological limitations have prevented a comprehensive understanding of DA release dynamics relative to reward. Many studies have primarily focused on transient changes in extracellular DA over seconds to minutes, which has limited our understanding of how slower, long-term changes in spontaneous DA release contribute to reward behavior. Therefore, studying circadian changes in DA may offer new insights into how reward behavior is modulated on a circadian timescale.

To determine how circadian time shapes DA signaling across longer timescales, we used long-term fiber photometry in freely behaving mice to measure both spontaneous and reward-evoked DA release. By pairing these recordings with an automated reward delivery system, we examined how DA dynamics and reward behavior evolve across the circadian cycle. We find that spontaneous and reward-evoked DA release exhibit distinct circadian rhythms and that their relationship to reward behavior depends on circadian time. Together, these findings suggest that the circadian system dynamically organizes multiple modes of DA signaling to shape reward processing across the day.

## RESULTS

### Spontaneous DA release in the NAc exhibits circadian rhythmicity

To measure spontaneous DA release across the circadian cycle, we performed long-term fiber photometry in freely behaving mice expressing the genetically-encoded dopamine sensor DA3m in the NAc. We injected mice with AAV9-hSyn-DA3m, implanted them with an optical fiber targeting the NAc, and recorded them for 10 min every 30 min across multiple days in a 12 h:12 h light:dark cycle (LD) followed by constant darkness (DD; **Fig. 1a**). Histological analysis confirmed DA3m expression and fiber placement in the NAc (**Fig. 1b**).

**Figure 1.**
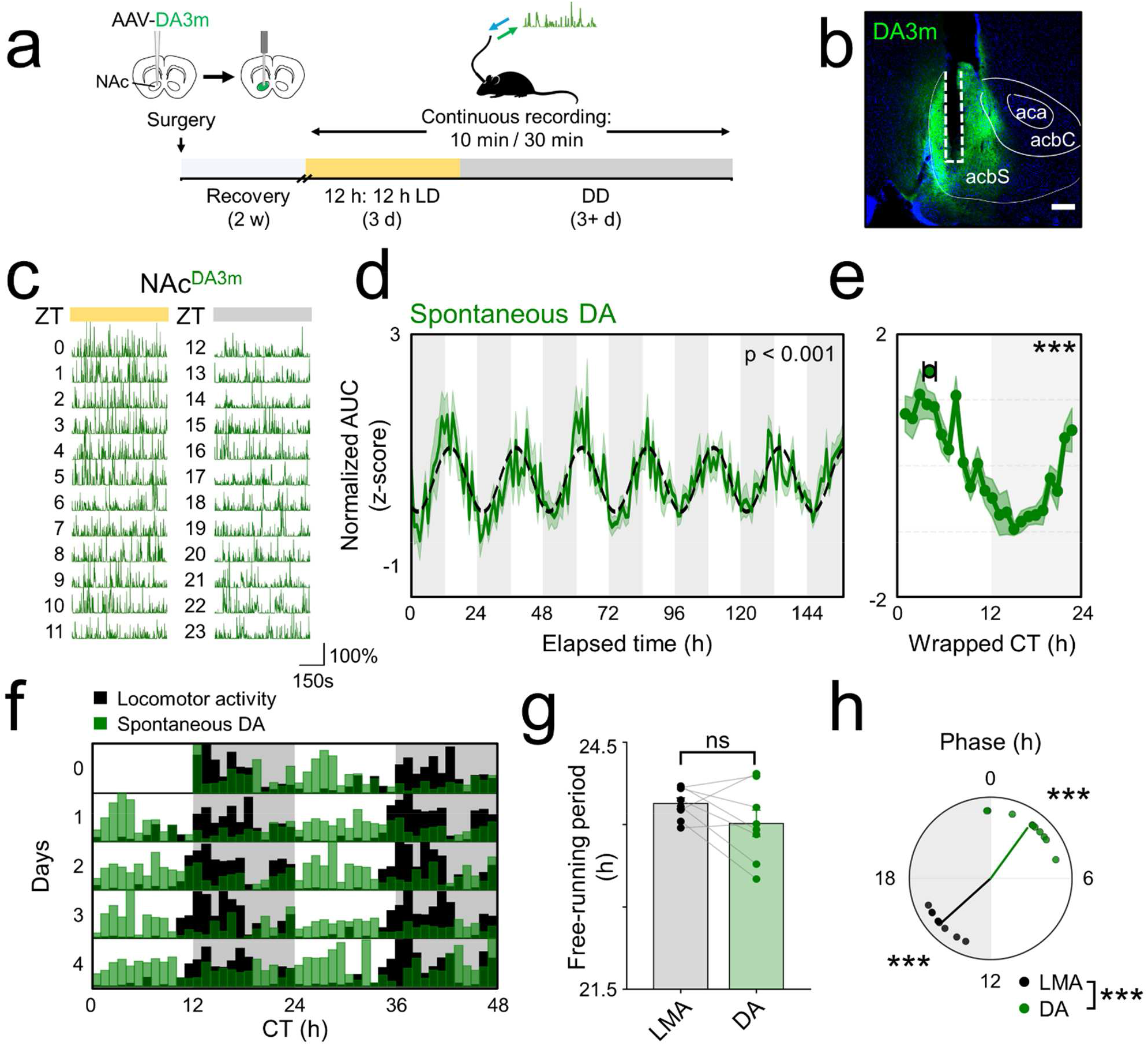
Spontaneous DA release in the NAc peaks opposite to locomotor activity. **a)** Schematic of the experimental design. AAV-DA3m was stereotaxically injected into the nucleus accumbens (NAc), followed by fiber optic implantation. After 2 w of recovery, spontaneous dopamine release was monitored by long-term fiber photometry in freely behaving mice, with 10 min recordings collected every 30 min for 3 d in a 12 h:12 h light:dark (LD) cycle followed by 3+ d in constant darkness (DD). **b)** Representative coronal section showing DA3m expression in the NAc (green) and the fiber optic implant tract (white dashed line). acbS, nucleus accumbens shell; acbC, nucleus accumbens core; aca, anterior commissure. **c)** Representative 10 min traces of normalized fluorescence signal (ΔF/F) recorded each hour across a 24 h day in LD from an individual mouse. Time is shown in zeitgeber time (ZT), where ZT 0 is lights-on. **d)** Group average of spontaneous DA3m fluorescence (z-scored, normalized area under the curve [AUC]) across the full DD recording period (n = 10 animals), plotted in elapsed recording time. White shading, subjective day; gray shading, subjective night; black dashed line, Cosinor fit (p < 0.001). **e)** Same data as in **d**, replotted in circadian time (modulo free-running period) and wrapped into a 24 h rhythm profile. Green line and shading depict the hourly mean + SEM. Green circle depicts the weighted center-of-mass (CoM; mean + SEM; Rayleigh test, p < 0.001). **f)** Representative double-plotted actogram depicting spontaneous DA3m fluorescence (green) and locomotor activity (black) recorded from an individual mouse in DD. **g)** Within-animal free-running periods of locomotor activity (Lomb-Scargle periodogram; τ = 23.76 ± 0.07 h) and DA3m fluorescence (τ = 23.51 ± 0.16 h) were not significantly different (paired t-test, p > 0.05). **h)** Rayleigh plot showing peak phase of spontaneous DA3m fluorescence (green; CT 2.4 ± 1.58; R = 0.918) and locomotor activity (black; CT 15.20 ± 0.92; R = 0.972). Circles represent individual mice, and arrows indicate the Rayleigh mean vector. Peak phases differed significantly (Watson-Williams test, p < 0.001). Analyses in **g** and **h** included n = 8 animals with rhythmic DA3m fluorescence and usable wheel-running data. *, p < 0.05; **, p < 0.01; ***, p < 0.001.

Representative ΔF/F traces from each hour across a single LD cycle from an individual mouse qualitatively showed higher DA3m fluorescence during the day than during the night (**Fig. 1c**). To quantify this pattern, we calculated the area under the curve (AUC) for each ΔF/F trace. Across animals, (n = 10), spontaneous DA3m fluorescence exhibited a significant circadian rhythm in DD as determined by Cosinor analysis (p < 0.001; Fig. 1d). At the individual-animal level, 9 of 10 mice showed significant circadian rhythmicity by Lomb-Scargle periodogram analysis (p < 0.05). After converting each animal’s data to circadian time, we found that the weighted center of mass (CoM) for spontaneous DA3m fluorescence occurred at circadian time (CT) 3.4 ± 0.8 (**Fig. 1e**). EGFP-expressing control animals (n = 2) did not show significant circadian rhythmicity (**Supplementary Fig. 1**).

We next compared spontaneous DA3m rhythms with locomotor activity measured by wheel running. Whereas locomotor activity peaked during the subjective night, spontaneous DA3m fluorescence peaked during the subjective day (**Fig. 1f**). Free-running period did not differ significantly between locomotor activity and spontaneous DA3m fluorescence within animals (paired t-test, p > 0.05; **Fig. 1g**). Using Rayleigh mean vector analysis, we determined locomotor activity peaked at CT 15.20 ± 0.92, whereas spontaneous DA3m fluorescence peaked at CT 2.4 ± 1.58, and these peak phases differed significantly (Watson-Williams test, p < 0.001; **Fig. 1h**; within-animal actograms and Rayleigh plots are shown in **Supplementary Fig. 2**).

Together, these results show that spontaneous DA release in the NAc is circadian, peaks during the subjective day, and is opposite in phase to locomotor activity.

### Automated reward delivery enables simultaneous long-term measurement of reward behavior and DA release

We next asked whether reward-evoked DA release is similarly organized across circadian time. To test this, we developed an automated paradigm for hourly sucrose delivery with simultaneous NAc DA3m photometry recordings over multiple days in LD and DD (**Fig. 2a,b**). Following acclimation to 5% sucrose, mice were given 10 min/h access to sucrose through an automated sliding door while receiving ad libitum access to water. Photometry recordings began 5 min before reward availability and continued throughout the 10 min reward-access period, yielding 15 min of recording per hour (**Fig. 2b**). Custom lickometers recorded sucrose licks and water licks across the experiment (**Fig. 2c**). In individual trials, sucrose access was accompanied by a discrete increase in DA3m fluorescence, enabling computational identification of reward-evoked DA peaks (**Fig. 2d**; **Supplementary Fig. 3**). This approach enabled longitudinal extraction of multiple behavioral and DA-related variables across the circadian cycle, including sucrose licks, water licks, bout counts, response speed, total DA, and peak DA (**Fig. 2e**; **Supplementary Fig. 4**).

**Figure 2.**
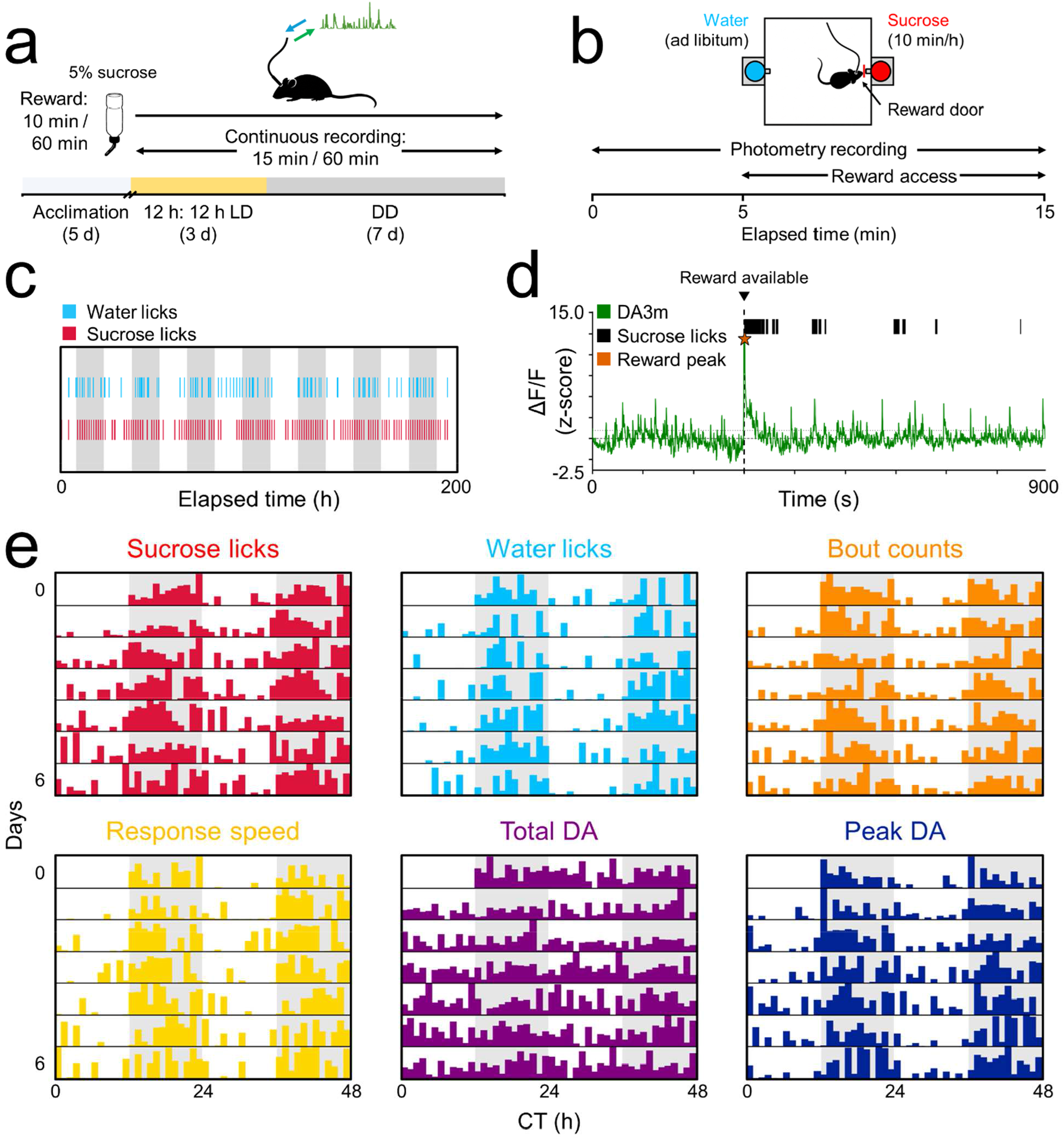
Automated reward delivery enables long-term simultaneous measurement of DA release and reward-related behavior. **a**,**b)** Schematic of the experimental design. Following acclimation to 5% sucrose for 5 d, DA3m-expressing mice underwent automated hourly sucrose presentation combined with long-term fiber photometry recording for 3 d in a 12 h:12 h light:dark (LD) cycle followed by 7+ d in constant darkness (DD). During each recording hour, photometry signals were recorded for 15 min, beginning 5 min before reward availability. Mice had ad libitum access to water and 10 min/h access to 5% sucrose through an automated reward door. **c)** Ethogram from an individual mouse showing water licks (blue) and sucrose licks (red) across the recording period in DD. White shading, subjective day; gray shading, subjective night. **d)** Representative 15 min photometry trace from an individual mouse aligned to reward availability. Green trace, DA3m fluorescence (ΔF/F); vertical dashed line, onset of reward availability; black tick marks, individual sucrose licks; orange star, automatically detected reward-evoked DA3m peak. **e)** Six representative double-plotted actograms from an individual mouse showing the range of behavioral and photometry-derived measures monitored continuously across multiple days, including sucrose licks, water licks, bout counts, response speed, total dopamine (DA), and peak DA. White shading, subjective day; gray shading, subjective night.

### Reward-related behaviors peak during subjective night

We first asked whether these reward-related behavioral variables were themselves organized across circadian time. To test this, we quantified circadian patterns of sucrose licks, water licks, bout counts, and response speed in DD (**Fig. 3a-d**). Response speed was evaluated only for time points in which sucrose licking occurred. Sucrose licks, bout counts, and response speed directly reflected behavior during reward access, whereas ad libitum water licking provided a non-reward control measured in the same animals. Across animals (n = 8), all four measures exhibited significant circadian rhythmicity by Cosinor analysis (p < 0.001 for all measures). After conversion to circadian time, all four behaviors peaked in the early subjective night by weighted CoM analysis. Wrapped circadian profiles plotted in raw units are shown in Supplementary Fig. 5. Consistent with this, individual-animal peak phases were significantly clustered for all four measures by Rayleigh analysis, again placing peak phase in the early subjective night. Together, these results show that multiple reward-related behaviors are circadian and peak during the early subjective night.

**Figure 3.**
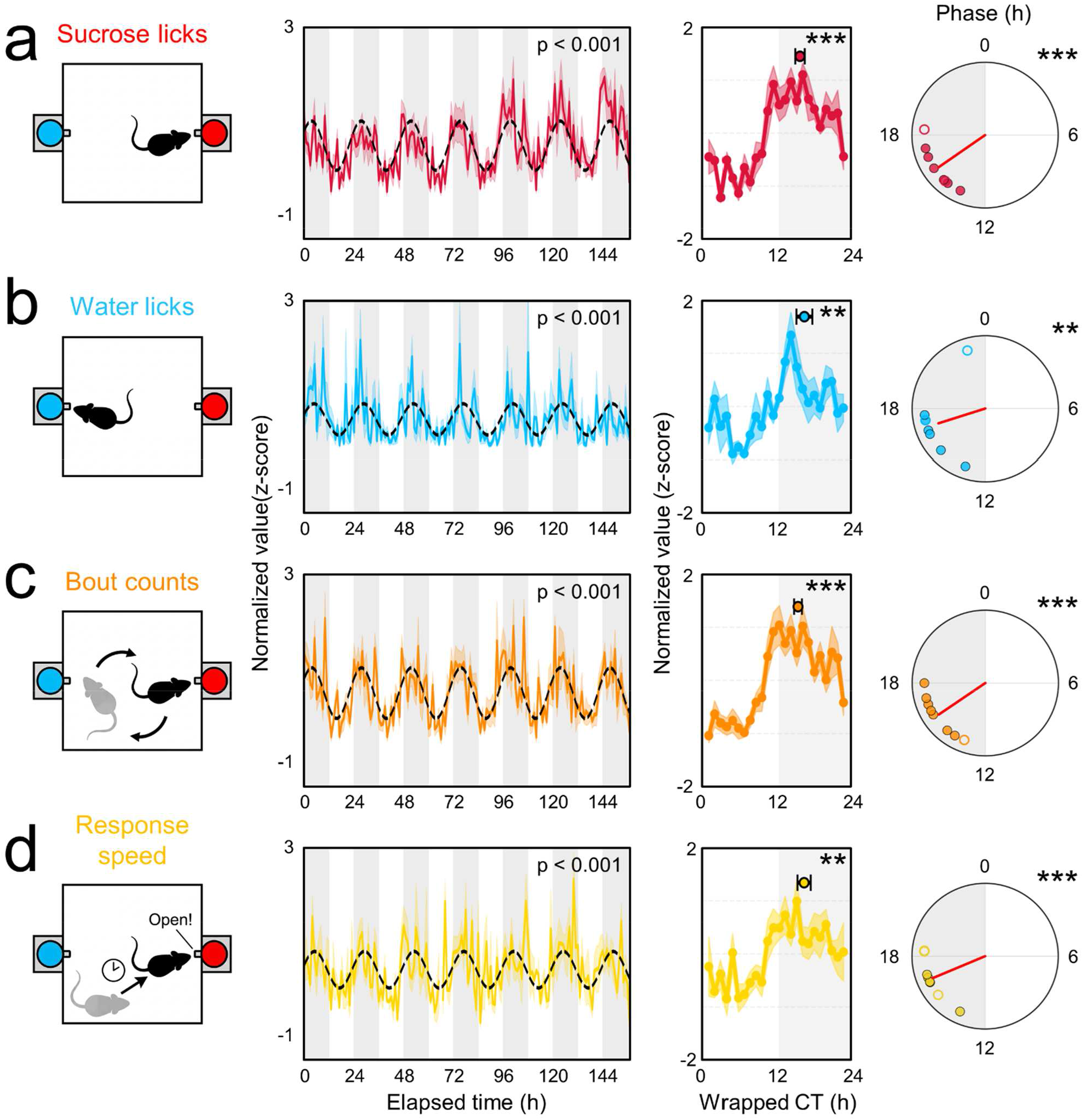
Reward behaviors peak during subjective night. **a-d)** Rows depict sucrose licks (red), water licks (blue), bout counts (orange), and response speed (yellow). In each row, the left panel is a schematic of the measured behavior. The middle-left panel shows the group average of the normalized, z-scored measure across the full constant darkness (DD) recording period, plotted in elapsed recording time. White shading, subjective day; gray shading, subjective night; black dashed line, Cosinor fit (p < 0.001 for all measures). The middle-right panel shows the same data replotted in circadian time (modulo free-running period) and wrapped into a 24 h rhythm profile. Line and shading depict the hourly mean ± SEM, and the circle indicates the weighted center of mass (CoM; n = 8 animals; Rayleigh test: sucrose licks, CT 15.5 ± 0.8, p < 0.001; water licks, CT 16.3 ± 1.4, p < 0.01; bout counts, CT 15.2 ± 0.7, p < 0.001; response speed, CT 16.2 ± 1.1, p < 0.01). The right panel shows Rayleigh plots of peak phase (n = 8 animals; Rayleigh test: sucrose licks, CT 15.7 ± 1.5, p < 0.001; water licks, CT 16.83 ± 2.5, p < 0.01; bout counts, CT 15.7 ± 1.5, p < 0.001; response speed, CT 16.5 ± 1.4, p < 0.001). Circles represent individual mice, and arrows indicate the Rayleigh mean vector. Filled circles denote individually rhythmic animals, and open circles denote individually arrhythmic animals (Lomb-Scargle periodogram; sucrose licks, 6/8 rhythmic; water licks, 6/8 rhythmic; bout counts, 7/8 rhythmic; response speed, 5/8 rhythmic). In **d)**, only event-locked trials were included in the middle two panels. *, p < 0.05; **, p < 0.01; ***, p < 0.001.

### Baseline, total, and reward-evoked peak DA show distinct circadian patterns

We next asked whether delineating the DA3m signal into pre-reward and reward-evoked components would alter its circadian profile. To test this, we quantified baseline DA, defined as the mean pre-reward DA3m signal; total DA, defined as post-reward area under the curve of the DA3m signal; and peak DA, defined as the maximum reward-evoked DA3m response (**Fig. 4a-c**). Because peak DA depends on reward engagement, group-level circadian analyses for this measure included only time points in which sucrose licking occurred.

**Figure 4.**
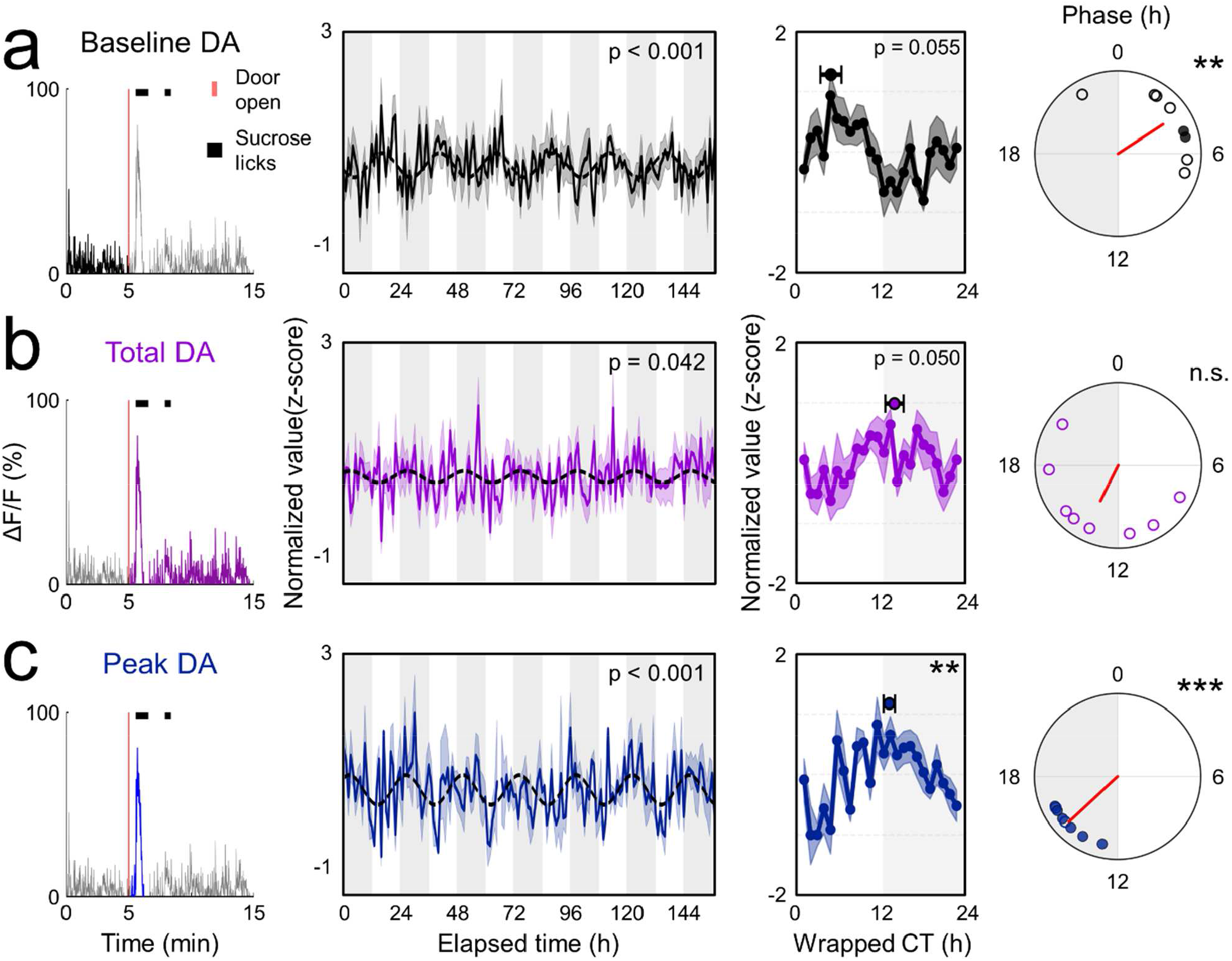
Baseline, total, and reward-evoked peak DA show distinct circadian patterns. **a-c)** Rows depict baseline DA, total DA, and peak DA, quantified from the DA3m photometry signal during the reward trial. In each row, the left panel depicts a representative DA3m photometry trace from an individual mouse during a 15 min reward trial. The red vertical line indicates reward availability, black tick marks indicate individual sucrose licks, and shaded regions indicate the portion of the trace used for quantification. Baseline DA (black) was defined as the mean DA3m signal during the 5 min period before reward availability. Total DA (purple) was defined as the area under the curve during the 10 min reward access period. Peak DA (blue) was defined as the maximum reward-evoked DA3m response during reward access. The middle-left panel shows the group average of the normalized, z-scored measure across the full constant darkness (DD) recording period, plotted in elapsed recording time. White shading, subjective day; gray shading, subjective night; black dashed line, Cosinor fit (baseline DA, p < 0.001; total DA, p = 0.042; peak DA, p < 0.001). The middle-right panel shows the same data replotted in circadian time (modulo free-running period) and wrapped into a 24 h rhythm profile. Line and shading depict the hourly mean ± SEM, and the circle indicates the weighted center of mass (CoM; n = 8 animals; Rayleigh test: baseline DA, CT 4.1 ± 1.6, p = 0.055; total DA, CT 13.6 ± 1.4, p = 0.050; peak DA, CT 12.8 ± 0.8, p < 0.01). The right panel shows Rayleigh plots of peak phase (n = 8 animals; Rayleigh test: baseline DA, CT 3.78 ± 2.72, p < 0.01; total DA, CT 13.77 ± 4.02, p > 0.05; peak DA, CT 15.14 ± 1.12, p < 0.001). Circles represent individual mice, and arrows indicate the Rayleigh mean vector. Filled circles denote individually rhythmic animals, and open circles denote individually arrhythmic animals (Lomb-Scargle periodogram; baseline DA, 2/8 rhythmic; total DA, 0/8 rhythmic; peak DA, 6/8 rhythmic). In **c)**, only event-locked trials were included in the middle two panels. *, p < 0.05; **, p < 0.01; ***, p < 0.001.

Across animals (n = 8), all three measures exhibited significant circadian rhythmicity by Cosinor analysis, although the profile of these rhythms differed (baseline DA, p < 0.001; total DA, p = 0.042; peak DA, p < 0.001). After conversion to circadian time, peak DA showed a significant weighted CoM that fell in the early subjective night, whereas the weighted CoMs for baseline DA and total DA were not significant. Consistent with this, Rayleigh analysis of individual-animal peak phases showed significant clustering for baseline DA in the early subjective day (p < 0.01) and for peak DA in the early subjective night (p < 0.001), but not for total DA (p > 0.05). Together, these results indicate that baseline and reward-evoked peak DA occupy distinct circadian phases, whereas total DA shows weaker rhythmic structure.

### Circadian phase-dependent coupling of peak DA to reward behavior and baseline DA

Finally, we asked whether peak DA tracked reward behavior and baseline DA differently across circadian time. To test this, we applied linear mixed models to hourly DD data from trials in which sucrose licking occurred, grouping observations into four circadian time bins (CT 0-6, 6-12, 12-18, and 18-24; **Fig. 5**).

**Figure 5.**
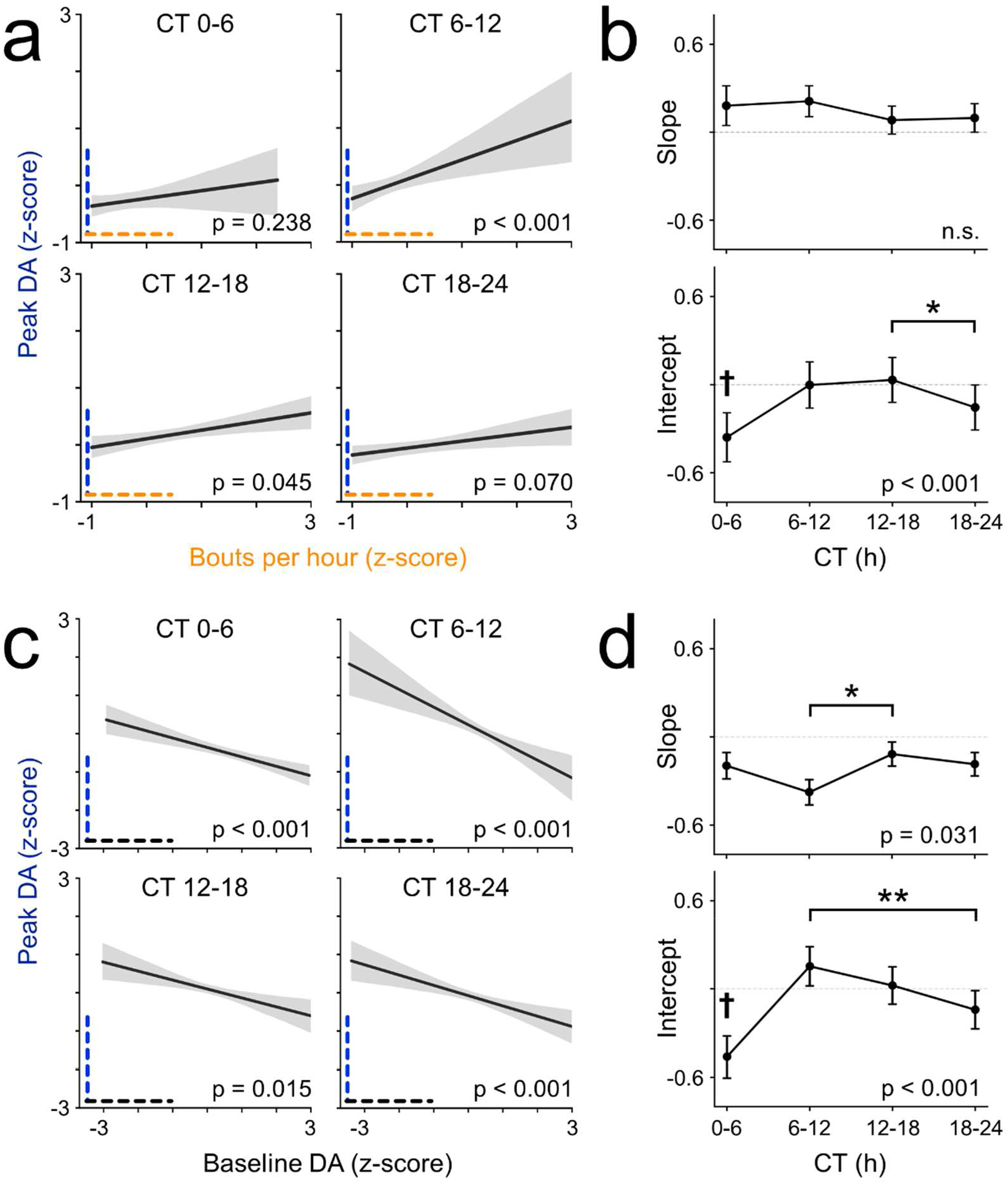
The relationship of peak DA to reward behavior and baseline DA depends on circadian phase. **a-d)** Linear mixed model (LMM) analyses were used to assess the relationship between bouts per hour and peak DA (**a**,**b**) or baseline DA and peak DA (**c**,**d**) across four circadian time bins (CT 0-6, 6-12, 12-18, and 18-24; n = 8 animals). In **a** and **c**, black lines indicate the fitted LMM relationship and gray shading indicates the 95% confidence interval. P values indicate whether the within-bin slope differed from zero using a Wald z-test. In **b** and **d**, slope (top) and intercept (bottom) estimates are plotted across circadian time bins for individual animals. Likelihood ratio tests were used to assess the effect of circadian time on slope and intercept, followed by post hoc Wald tests with Holm-Bonferroni correction. **a)** Relationship between bouts per hour and peak DA (CT 0-6, p = 0.238; CT 6-12, p < 0.001; CT 12-18, p = 0.045; CT 18-24, p = 0.070). **b)** Circadian effects on slope and intercept for the model in a (slope, p > 0.05; intercept, p < 0.001). No post hoc differences were detected for slope. For intercept, CT 12-18 and CT 18-24 differed (p < 0.05), and CT 0-6 differed from all other time bins (p < 0.05). **c)** Relationship between baseline DA and peak DA (CT 0-6, p < 0.001; CT 6-12, p < 0.001; CT 12-18, p = 0.015; CT 18-24, p < 0.001). **d)** Circadian effects on slope and intercept for the model in **c** (slope, p = 0.031; intercept, p < 0.001). For slope, CT 6-12 differed from CT 12-18 (p < 0.05). For intercept, CT 6-12 and CT 18-24 differed (p < 0.01), and CT 0-6 differed from all other time bins (p < 0.05). *, p < 0.05; **, p < 0.01; ***, p < 0.001; †, p < 0.05 versus all other time bins.

We first examined the relationship between peak DA and bouts per hour (**Fig. 5a,b**). Within-bin slopes were significantly different from zero during CT 6-12 and CT 12-18 (Wald z-test, p < 0.05), but not during CT 0-6 or CT 18-24. Circadian phase did not significantly alter the slope of this relationship (LRT, p > 0.05), indicating that the rate of change between bouts and peak DA was similar across time bins. However, circadian phase did affect the intercept, indicating a phase-dependent shift in the scale of peak DA relative to bout count(LRT, p < 0.05, Wald z-test, p < 0.05). Specifically, the intercept at CT 0-6 was lower than at all other time bins, and the intercept at CT 12-18 was higher than at CT 18-24.

We next examined the association between baseline DA and peak DA (**Fig. 5c,d**). Within-bin slopes were significantly different from zero in all four circadian time bins (Wald z-test, p < 0.05). In contrast to the bouts analysis, circadian phase significantly affected both slope and intercept (LRT, p < 0.05, Wald z-test, p < 0.05), indicating that circadian phase altered both the rate of change and offset between baseline DA and peak DA. Specifically, the slope differed between CT 6-12 and CT 12-18. For intercept, CT 0-6 was lower than all other time bins, and CT 6-12 differed from CT 18-24.

Together, these results show that the circadian phase differentially shapes the relationship of peak DA to reward behavior and baseline DA, shifting the reward-behavior relationship primarily through changes in intercept and the baseline-peak DA relationship through changes in both slope and intercept.

## DISCUSSION

Previous studies have established that the circadian system is an important modulator of DA dynamics and reward behavior [18–24]. Similarly, mesolimbic DA activity is a well-established neural correlate of reward behavior [15,25,26]. However, how the circadian system modulates this association has remained unclear. In this study, we directly probed the DA-reward axis on a circadian timescale using long-term in vivo imaging paired with automated reward delivery. Our results revealed distinct circadian rhythms in spontaneous and reward-evoked DA release. We also found that the relationship between reward-evoked peak DA and reward behavior depended on circadian time, and that the relationship between baseline and reward-evoked DA was likewise subject to circadian modulation. Altogether, these findings support a model in which circadian time dynamically tunes DA release and its relationship to reward behavior.

The study of how DA release changes over time *in vivo* has been historically challenging. Although prior studies using microdialysis and fast-scan cyclic voltammetry have provided important insights, limitations in temporal resolution or longitudinal stability have made it difficult to examine DA release continuously across circadian timescales in vivo [17,23,24,27]. Here, we addressed these limitations using long-term fiber photometry, which enabled specific, sub-second detection of DA-related signals over several days. In our first experiment, we sought to identify broad changes in DA by measuring circadian rhythms in spontaneous DA release. Based on prior microdialysis studies reporting elevated striatal DA at night, we initially expected spontaneous DA release to peak during the active phase. However, we instead observed a circadian rhythm in spontaneous DA release that peaked early in the subjective day. Importantly, the average free-running period of spontaneous DA release was less than 24 h and did not differ from the average wheel-running free-running period, supporting the use of long-term photometry for circadian measurements. One possible explanation for this discrepancy is that DA3m fluorescence reflects instantaneous DA bioavailability rather than absolute extracellular DA concentration, and may therefore be differentially influenced by factors such as uptake kinetics. Consistent with this interpretation, evoked DA release in the NAc has been reported to be greater during the day under tonic stimulation conditions, suggesting enhanced sensitivity to low-frequency or spontaneous DA signaling during this phase [20]. Alternatively, the daytime peak in spontaneous DA release may reflect a biologically distinct state-dependent process, such as changes in VTA dopaminergic neuron activity associated with sleep [28].

These findings prompted us to examine how rhythms in spontaneous DA release relate to reward-related behavior. In our second experiment, we used long-term fiber photometry paired with an automated reward delivery system to measure DA release and reward behavior simultaneously. We found that 5% sucrose was sufficient to elicit a detectable spike in DA release. As expected, all measured reward behaviors exhibited circadian rhythms that peaked in early subjective night at both the individual and group levels. These findings align with prior studies showing greater natural reward engagement at night.[22,29] However, to our knowledge, this is among the first studies to track circadian rhythms in multiple reward-related behaviors across several days while maintaining high temporal resolution. Importantly, mice continued to engage robustly with the sucrose reward despite repeated exposures, indicating stable reward-directed behavior throughout the experiment. Because mice had ad libitum access to food and water, this engagement was unlikely to be driven solely by hunger or thirst.

To gain a clearer understanding of how DA release changed throughout the reward experiment, we delineated the DA3m signal into three modes: baseline, total, and peak DA. Baseline DA in our second experiment reflects spontaneous DA activity outside of reward-evoked transients and is therefore directly comparable to spontaneous DA measured in our first experiment. Consistent with that comparison, baseline DA showed a peak phase near CT 4 by Rayleigh analysis, despite otherwise exhibiting weaker rhythmicity. Total DA exhibited the weakest rhythmic profile, reaching significance at the group level but not in individual-animal phase analyses. Peak DA, however, was rhythmic at both the group and individual-animal levels, indicating that the transient reward-evoked component of the DA signal carries clear circadian structure that is otherwise masked by the total DA signal measured across the full 10 min reward period.

Center-of-mass and Rayleigh analyses provided complementary views of peak DA rhythmicity. Using these two approaches, we found that both the frequency and magnitude of peak DA varied across circadian time, respectively. Rayleigh analysis showed that the frequency of peak DA events closely aligned with the phase of reward behavior, indicating that this metric more closely reflects event-locked DA responses. In contrast, center-of-mass analysis showed that peak DA magnitude peaked near the subjective day-to-night transition (CT 12.8), slightly earlier than the peak in reward behavior (CT 15.5). Together, these findings indicate that the circadian modulation of peak DA is not fully explained by the timing of reward behavior alone. More broadly, they suggest that circadian modulation of DA release is multimodal, with distinct rhythms in baseline and peak DA signals. Although the mechanisms underlying these two modes remain unclear, prior work has shown that cholinergic inputs in the NAc and dopaminergic neurons in the VTA differentially regulate DA release, suggesting a potential circuit-level basis for the distinct rhythmic patterns observed here [20].

Our final line of investigation was to more precisely characterize the relationships between peak DA, reward behavior, and baseline DA. Using LMMs, we found that the relationship between peak DA and reward behavior (bouts per hour) was significant only during the late subjective day and early subjective night. While the slope remained stable across circadian time, the intercept varied, mirroring the phase structure we observed for peak DA in our CoM analysis. This indicates that equivalent levels of reward behavior were associated with different magnitudes of peak DA depending on circadian phase. In addition, baseline and peak DA were negatively correlated across all circadian phases, suggesting a balance between spontaneous and reward-evoked DA signaling whereby lower baseline DA was associated with greater reward-evoked responses. One possible interpretation is that this relationship reflects variation in DA bioavailability, such that reduced spontaneous release may give way to greater DA availability for presynaptic exocytosis during reward. Consistent with this possibility, prior studies have shown that presynaptic DA availability varies with time of day [27]. We also found that the slope and intercept of the baseline-peak DA relationship changed across circadian phase, with the most pronounced shift occurring during the late subjective day. Together, these findings suggest that circadian modulation of the baseline-peak DA axis may shift DA signaling towards a reward-directed state during the day-to-night transition.

Our collective results reveal a complex interplay between the circadian system and mesolimbic DA release. Spontaneous DA release peaked during the early subjective day, reward-evoked DA release peaked near the day-to-night transition, and reward behavior peaked during early subjective night. We further found that circadian time is an important determinant of how DA release relates to behavior and how spontaneous and reward-evoked DA relate to each other. Ultimately, these results support a model in which circadian modulation of baseline DA may alter the gain of reward-evoked signaling, amplifying DA responses during behaviorally relevant times of day. More broadly, these findings suggest that circadian state is an important and underappreciated determinant of mesolimbic DA function, with implications for how reward-related neural signals are encoded across time of day.

## METHODS

### Animals and housing

Male and female wild-type mice (2 to 4 months old) on a C57BL/6J background were used for all experiments. Mice were singly housed at constant temperature (∼23°C) and humidity (∼40%) under a 12 h:12 h light:dark cycle (LD; lights on defined as zeitgeber time 0) for 3 d, followed by 7 d in constant darkness (DD), with food and water available ad libitum. Nesting material and bedding were provided in all home cages, and running wheels were included except during experiments using the reward behavior paradigm (see below). All experiments were approved by and performed in accordance with the guidelines of Texas A&M University’s Institutional Animal Care and Use Committee.

### Stereotaxic surgery

Mice were anesthetized using a SomnoSuite low-flow anesthesia system (Kent Scientific) with induction at 4% isoflurane, 400 mL/min and maintenance at 2% isoflurane, 30 mL/min throughout surgery. Buprenorphine-SR (1 mg/kg) was administered subcutaneously before surgery. After the skull was leveled, AAV9-hSyn-DA3m (∼5.40 × 10^12^ GC/mL; BrainVTA) or AAV9-hSyn-EGFP (∼2.90 × 10^13^ GC/mL; University of Zurich Viral Vector Facility) [30] was diluted 1:4 in phosphate-buffered saline and injected unilaterally into the nucleus accumbens (NAc; AP +1.42 mm, ML +0.75 mm, DV -4.70 mm from bregma) at a volume of 200 nL and a rate of 1 nL/min. The injection needle was left in place for 10 min before withdrawal. A fiber optic cannula (6 mm in length, 200 µm diameter core, 0.39 NA; RWD Life Science) was implanted at the same coordinates and secured with a layer of adhesive cement (Parkell) and opaque black dental cement. The implant was covered with a thin layer of black nail polish. Mice were allowed to recover in their home cages for 2 weeks before experiments.

### Fiber photometry

Fiber photometry data were collected using a multi-channel fiber photometry system (FP3002; Neurophotometrics). After postsurgical recovery, mice were tethered to the photometry system and allowed at least 2 d to acclimate to the patch cable (200 µm diameter core, 0.37 NA, 3 m length; Doric Lenses) before the start of recordings. For spontaneous DA recordings, fluorescence was sampled at 40 frames/s using interleaved 470 nm (signal) and 415 nm (isosbestic) excitation channels using a 33% duty cycle (10 min on, 20 min off). For reward-evoked DA recordings, the fiber photometry system was triggered by a custom Arduino-based controller (TageLabs) to begin acquisition 5 min before reward availability and continue throughout the 10 min reward-access period, yielding 15 min of recording per hour (25% duty cycle). Reward-evoked recordings were also acquired at 40 frames/s using interleaved 470 nm and 415 nm excitation channels.

### Reward behavior paradigm

For long-term reward behavior experiments, mice were housed in custom-modified home cages fitted with two 3D-printed replacement walls. One wall provided access to a drinking spout containing water, and the other provided access to a drinking spout containing 5% sucrose solution as the reward stimulus. Both spouts were positioned behind infrared beam sensors that detected individual licks. Water was available ad libitum, whereas access to the 5% sucrose spout was restricted to 10 min once per hour using a servo-operated sliding door. Timing of reward access was controlled by a custom Arduino-based system, which triggered photometry acquisition each hour, opened and closed the reward door, and recorded infrared beam breaks with timestamps to a .csv file. The water-access wall was instrumented in the same manner, but without a sliding door. For acclimation, mice were first given water through both spouts for 2 d while the 10 min/h access schedule was active on one side. Mice then received water only through the gated spout for an additional 2 d before the start of the experiment. Successful licking from both spouts was confirmed during acclimation. Experiments consisted of 3 d in a 12 h:12 h light:dark cycle (LD) followed by 6 to 8 d in constant darkness (DD).

### Tissue collection and histology

At the conclusion of each experiment, mice were returned to a 12 h:12 h light:dark cycle for at least 3 d and then euthanized by transcardial perfusion at ZT 6. Mice were deeply anesthetized with isoflurane and transcardially perfused with phosphate-buffered saline followed by 4% paraformaldehyde. Brains were extracted, post-fixed overnight in 4% PFA, cryoprotected in 30% sucrose for 48 h, frozen in OCT compound, and sectioned coronally at 40 µm on a cryostat. Sections containing the NAc were mounted on slides and imaged on a Nikon Ti2-E equipped with an X-Light V3 spinning-disk confocal (Crest Optics) to verify DA3m expression and fiber tract location.

### Fiber photometry analysis

Raw fiber photometry data were processed using custom MATLAB and Python scripts based on the analysis framework described in [31]. To calculate ΔF/F, the 470 nm signal channel and 415 nm isosbestic channel were first low-pass filtered. A rolling ordinary least squares regression was then used to fit the isosbestic channel to the signal channel in 120 s windows while accounting for slow baseline drift. The fitted isosbestic signal was subtracted from the 470 nm signal to generate a drift-corrected fluorescence signal, which was then normalized to ΔF/F as previously described [31].

For spontaneous DA3m and EGFP recordings, area under the curve (AUC) was calculated for each 10 min ΔF/F trace, and these values were used for circadian analyses. For reward-evoked DA recordings, three values were extracted from each ΔF/F trace: the mean value during the 5 min pre-reward period (baseline DA), the area under the curve during the 10 min reward-access period (total DA), and the maximum value of the identified reward peak (peak DA). For total DA and peak DA analyses, traces were z-scored to the mean and standard deviation of the pre-reward baseline period before extracting the AUC and peak value, respectively.

Reward-evoked DA peaks were identified using the following pipeline. First, a continuous wavelet transform was applied to each z-scored ΔF/F trace. Wavelet magnitudes were then summed across frequency bands. Candidate reward-peak regions were defined as time periods in which the summed magnitude was greater than or equal to the 80th percentile of the trace. Candidate regions were then filtered behaviorally, retaining only those occurring within a valid sucrose-lick window extending from 10 s before to 1 s after an individual sucrose lick. The region with the highest summed magnitude within a valid lick window was selected, and the reward peak was defined as the maximum ΔF/F value within a 10 s window surrounding that region. As a behavioral failsafe, if sucrose licking occurred but no region exceeded the 80th percentile threshold, the threshold requirement was dropped and the highest summed-magnitude region within valid lick windows was used for reward peak detection.

### Behavior data analysis

For the spontaneous DA experiment, wheel-running data were recorded using ClockLab software (Actimetrics) and analyzed using custom Python scripts. Wheel-running activity was binned in 30 min intervals to match the sampling frequency of the spontaneous DA photometry data for pairwise comparisons.

For reward experiments, lickometer output was binary, with a value of 1 recorded when the infrared beam was broken and a value of 0 when the beam was restored. Each event was timestamped using Unix time. Because mice lick at a frequency of approximately 7 Hz [32]. lickometer events were filtered using a valid contact window of 50 to 100 ms to remove likely artifacts. Hourly lick counts were calculated from the number of valid beam-break events. For sucrose licking, hourly bout counts and response speed were also calculated. A bout was defined as a cluster of licks separated by less than 10 s, with pauses of 10 s or more defining a new bout. Response speed was calculated as the inverse of latency to first lick (s^-1^). To avoid division by zero, latency values were constrained to a minimum of 0.1 s.

### Statistical analysis

Statistical analyses were performed using custom Python scripts. For identification of circadian rhythmicity within individual animals, data from the first 7 d in constant darkness (DD) were analyzed using Lomb-Scargle periodogram analysis, with significance defined by false-alarm probability (p < 0.05) for periods between 22 and 26 h. Free-running periods for spontaneous DA and locomotor activity were compared using paired t-tests. Circular phase clustering and peak phase within individual animals were assessed using Rayleigh tests. For group-level analyses, peak phases from individual animals were aggregated and evaluated using Rayleigh tests, and group-level phase differences between spontaneous DA and locomotor activity were tested using Watson-Williams tests. Animals were included in these group-level circular analyses regardless of individual Lomb-Scargle significance, because individual rhythmicity and group-level phase clustering test different questions, and restricting the analysis to individually rhythmic animals would bias group estimates toward the strongest rhythms.

Additional group-level rhythmic analyses included Cosinor fitting and center of mass (CoM) calculations. For reward-dependent measures, including peak DA and response speed, only time points during which sucrose licking occurred were included for analysis. For Cosinor analysis, data from each animal across 7 d in DD were z-scored, averaged, and fit with a Cosinor model. For CoM analysis, 7 d of DD data from each animal were converted to circadian time, folded into a 24 h average profile, z-score normalized, and used to calculate individual CoMs, which were then averaged across animals using circular statistics.

To assess phase-dependent relationships between peak DA and bouts per hour or baseline DA, linear mixed-effects models (LMMs) were applied across four 6 h circadian time bins. These analyses included only trials in which sucrose licking occurred. Animal identity was included as a random intercept to account for repeated measures. Within-bin slope significance was assessed using Wald z-tests. To compare slopes and intercepts across circadian time bins, a global LMM including interactions between predictor and circadian time bin was evaluated using omnibus likelihood ratio tests, followed by post hoc Wald tests with Holm-Bonferroni correction where appropriate. Global z-scoring was used for LMM visualization only and did not alter statistical conclusions, which were verified using non-standardized data.

## CODE AVAILABILITY

Code used for analysis is available at https://github.com/jones-lab-tamu/DA_Photometry_and_Reward_Analysis

## ACKNOWLEDGMENTS

This work was supported by National Institutes of Health Grant R35GM151020 (J.R.J.), a research grant from the Whitehall Foundation (J.R.J.), and National Institutes of Health Grant F31DA065512 (J.N.C.). We thank members of the Jones Lab for helpful discussions and feedback on the manuscript.

## Notes

### Competing Interest Statement

The authors have declared no competing interest.

