## Supplementary Figures for "Circadian modulation of spontaneous dopamine release shapes reward-evoked signaling in the nucleus accumbens"

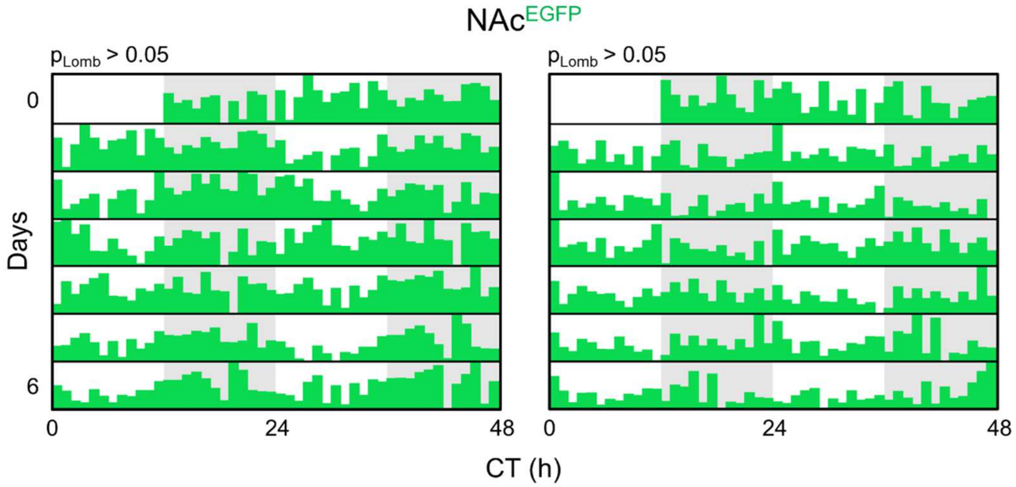


**Supplementary Figure 1. Photometry recordings of EGFP in the NAc do not exhibit significant circadian rhythmicity.** Two representative animals expressing EGFP in the NAc were recorded using long-term fiber photometry for 7 d in constant darkness (DD). Fluorescence was recorded for 10 min every 20 min. For each recording, ΔF/F was summarized as area under the curve (AUC) and plotted as a double-plotted actogram. Lomb-Ssargle periodogram analysis did not detect significant circadian rhythmicity in EGFP fluorescence in either animal ($p > 0.05$).

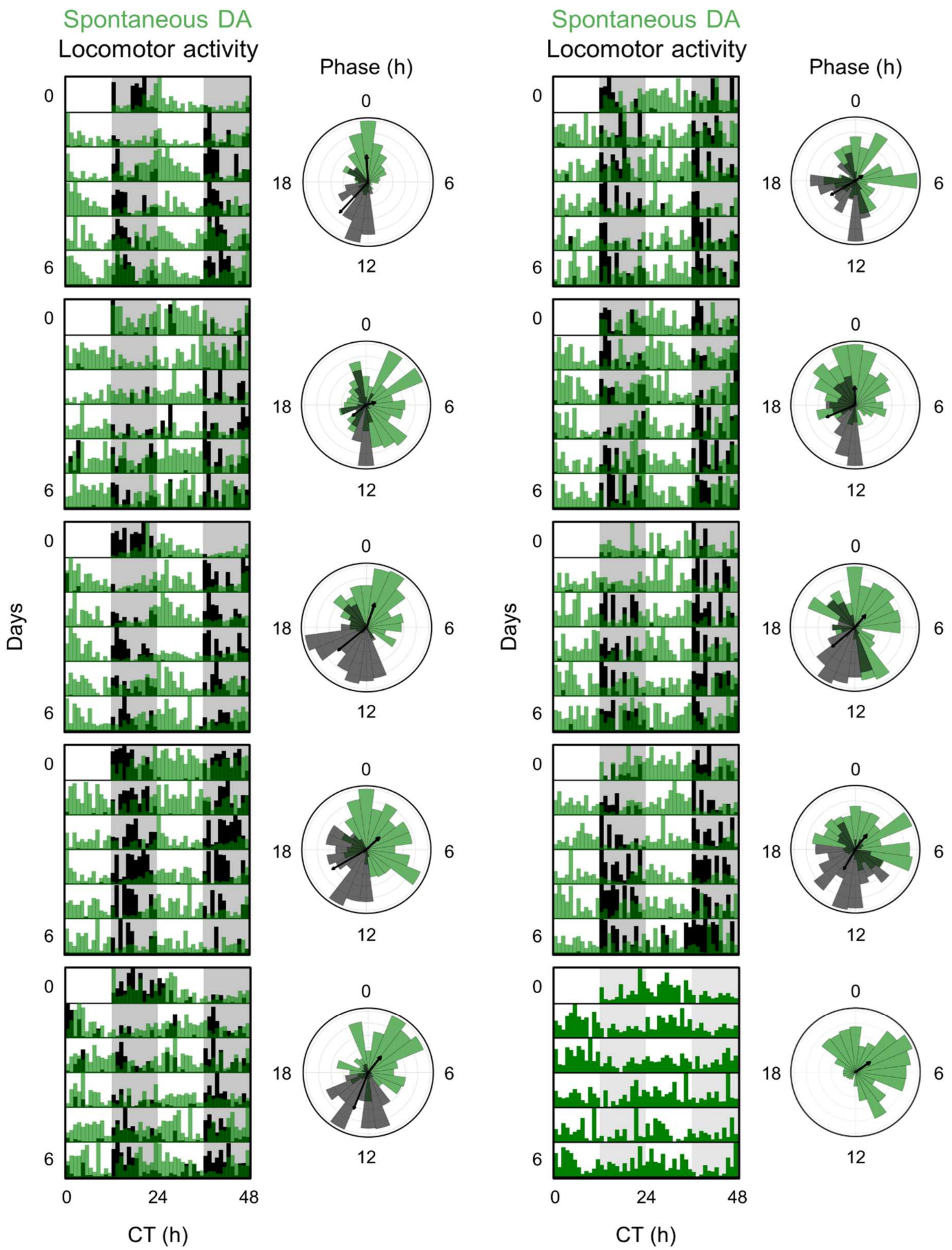


**Supplementary Figure 2. Spontaneous DA3m fluorescence peaks opposite to locomotor activity within individual animals.** Double-plotted actograms and corresponding Rayleigh plots from 10 individual animals showing spontaneous DA3m fluorescence (green) and locomotor activity (black) across 7 d in constant darkness (DD). Rayleigh plots depict repeated within-animal measurements collected every 30 min (2 data points per hour) across the recording interval. One animal (bottom right) did not have accompanying locomotor activity data and is therefore shown with spontaneous DA3m fluorescence only.

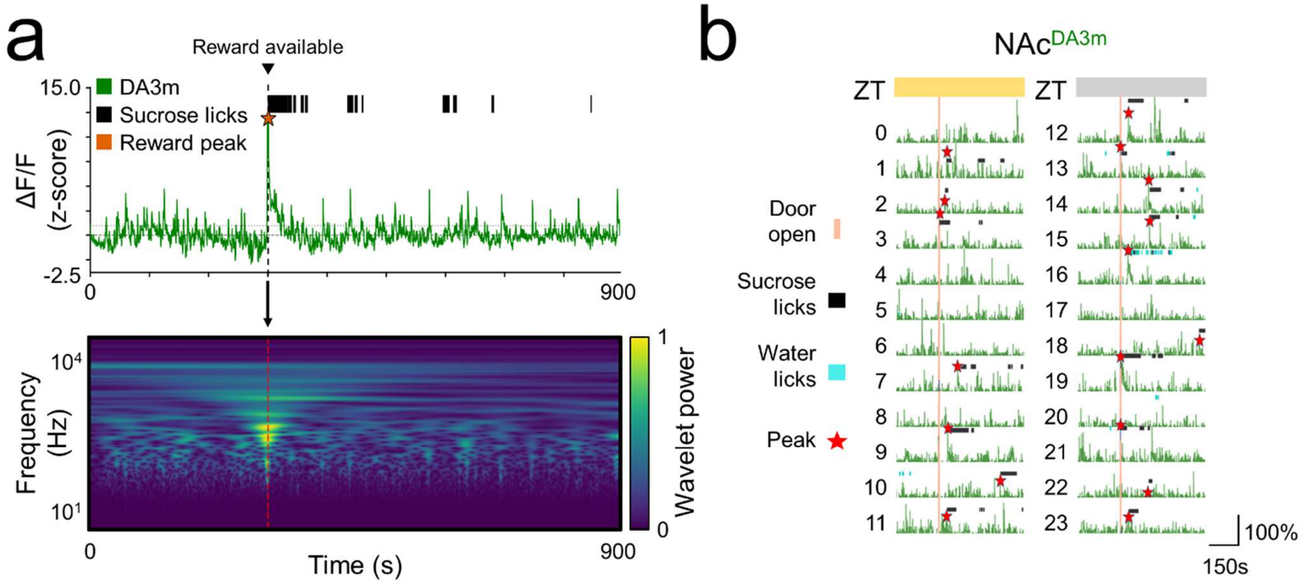


**Supplementary Figure 3. Reward-evoked DA peak identification by wavelet analysis. a)** Top, representative 15 min DA3m photometry trace from an individual mouse during a single reward trial. The vertical dashed line indicates reward availability, black tick marks indicate individual sucrose licks, and the red star indicates the automatically identified reward-evoked DA peak. Bottom, corresponding wavelet analysis of the same trace, plotted across time and frequency, with wavelet power shown by color scale. The time of maximal wavelet power aligns with the identified reward-evoked DA peak. **b)** Representative stacked 15 min DA3m photometry traces from each recording hour across a single day in a 12 h:12 h light:dark (LD) cycle. Time is shown in zeitgeber time (ZT), where ZT 0 is lights-on. The orange vertical line indicates reward availability, black tick marks indicate sucrose licks, blue tick marks indicate water licks, and red stars indicate automatically identified reward-evoked DA peaks.

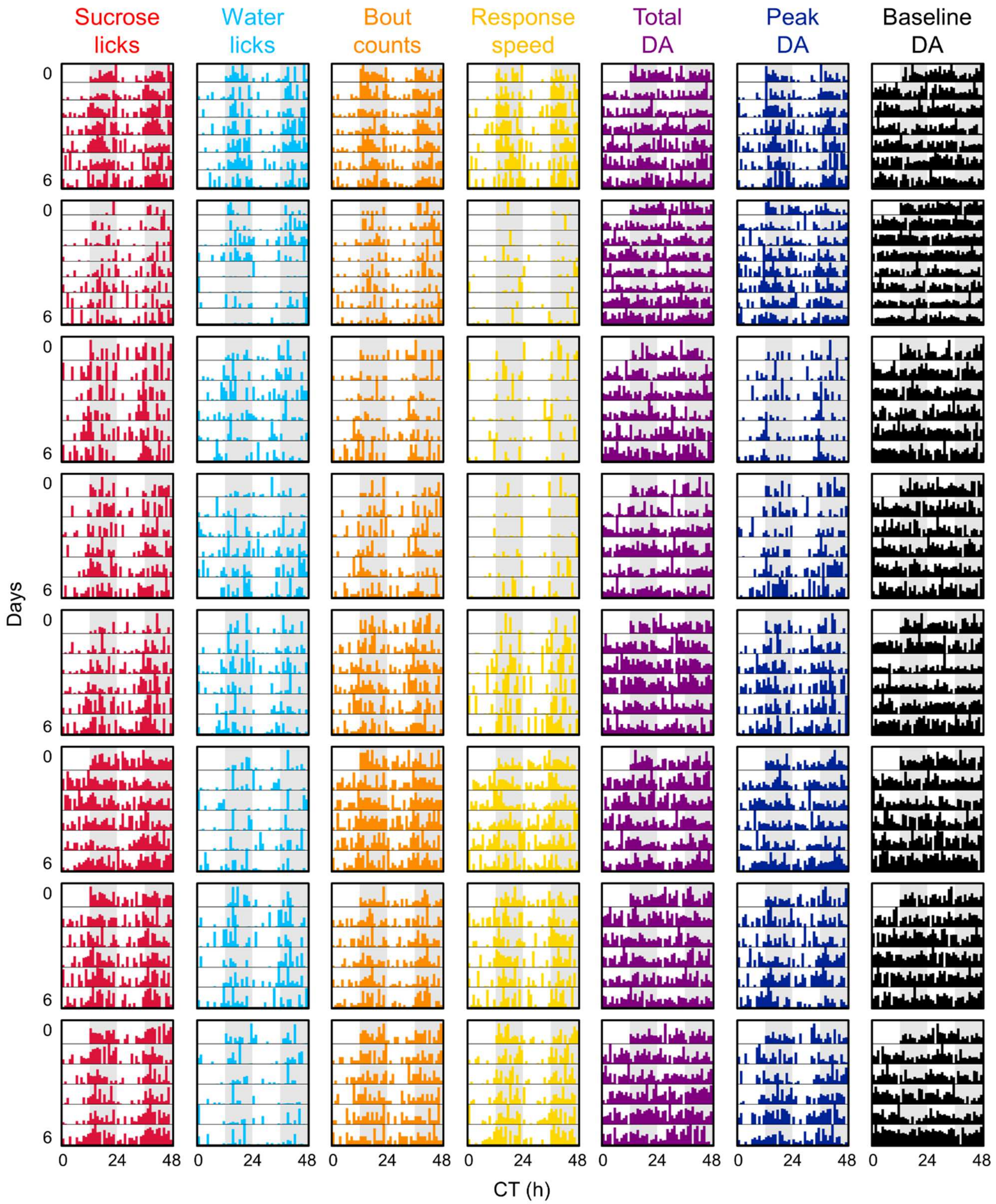


**Supplementary Figure 4. Individual-animal actograms for all measures during the reward experiment.** Each row shows double-plotted actograms from one of 8 individual animals across 7 d in constant darkness (DD). From left to right, panels show sucrose licks, water licks, bout counts, response speed, total DA, peak DA, and baseline DA.

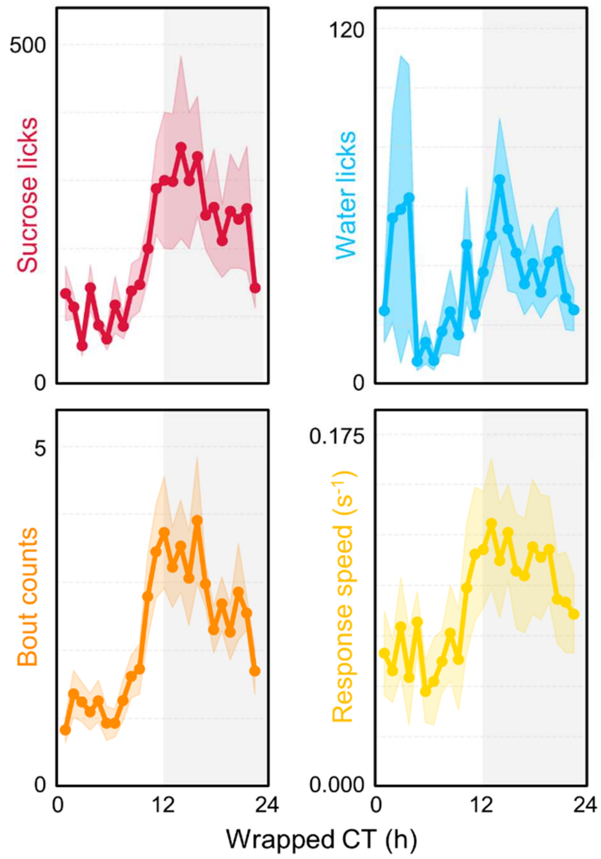


**Supplementary Figure 5. Wrapped circadian profiles of raw behavioral measures during the reward experiment.** Each plot shows the same circadian time-wrapped data presented in **Figure 3**, but using raw rather than normalized, z-scored values. Data were first averaged by circadian time within each mouse and then averaged across animals (n = 8). Lines and shading depict mean ± SEM. Measures shown are sucrose licks (red), water counts (blue), bout counts (orange), and response speed (yellow).
